# Visual signal evolution along complementary color axes in four bird lineages

**DOI:** 10.1101/514778

**Authors:** Anand Krishnan, Avehi Singh, Krishnapriya Tamma

## Abstract

Animal color patterns function in varied behavioral contexts including recognition, camouflage and even thermoregulation. The diversity of visual signals may be constrained by various factors, for example, dietary factors, and the composition of ambient environmental light (sensory drive). How have high-contrast and diverse signals evolved within these constraints? In four bird lineages, we present evidence that plumage colors cluster along a line in tetrachromatic color space. Additionally, we present evidence that this line represents complementary colors, which are defined as opposite sides of a line passing through the achromatic point (putatively for higher chromatic contrast). Finally, we present evidence that interspecific color variation over at least some regions of the body is not constrained by phylogenetic relatedness. Thus, we hypothesize that species-specific plumage patterns within these bird lineages evolve by swapping the distributions of a complementary color pair (or dark and light patches in one group, putatively representing an achromatic complementary axis). The relative role of chromatic and achromatic contrasts in discrimination may depend on the environment that each species inhabits.

## Introduction

The diverse colors of birds are important communication signals for advertisement and species recognition (Baker and Parker 1979; Alatalo et al. 1994; Greene et al. 2000; Bleiweiss 2004; Uy et al. 2009; Seddon et al. 2013). A number of ecological factors may constrain the diversification of plumage color, including diet (Hill et al. 2002; McGraw and Nogare 2004), the sensory systems of species and their predators(Gotmark 1993; Marchetti 1993; Gomez and Théry 2007), the composition of ambient light in different habitats (Endler 1992; Boughman 2002), and the additional constraint of phylogenetic relatedness (the tendency of related species to resemble each other). These constraints may operate both on what colors a bird may exhibit, and in where they are placed on the body (Endler 1992; Gomez and Théry 2007). Multiple studies have shown that plumage (and egg) colors in birds do not occupy the entire available color space (Endler et al. 2005; Cassey et al. 2008; Stoddard and Prum 2011). This constrained diversity of color has important consequences for signal perception and discrimination by receiver visual systems (Endler 1992; Endler and Mielke 2005; Cole and Endler 2016). In spite of these constraints, many bird groups have evolved highly diverse colors and patterns (Hill and McGraw 2006a); understanding signal evolution remains a focus of research (Stoddard and Prum 2008; Mason et al. 2014; Doutrelant et al. 2016).

The evolution of color signals may be influenced by how they are perceived by receivers, and this perception involves two neural processes: 1) photoreceptors (cones) sensitive to different wavelengths of light, and 2) downstream neural mechanisms comparing photoreceptor output, both spectrally and spatially (Endler and Mielke 2005; Endler et al. 2005; Kelber 2016). Neurons performing these comparisons possess a distinct spatial receptive field(Ventura et al. 2001), and thus likely respond differently to different patterns of the same pair of colors. Birds possess a tetrachromatic visual system, with ultraviolet-sensitive UVS (or violet-sensitive VS)(Ödeen and Håstad 2013), short (blue, SWS), medium (green, MWS) and long-wavelength (red, LWS) sensitive cones(Vorobyev et al. 1998; Vorobyev 2003; Osorio and Vorobyev 2008). Opponent-color mechanisms in birds support color discrimination, and multiple studies have investigated the behavioral abilities of birds to discriminate colors (Yazulla and Granda 1973; Wright 1975; Goldsmith and Goldsmith 1979; Osorio et al. 1999a,b; Vorobyev 2003; Goldsmith and Butler 2005; Ham and Osorio 2007).

How do visual signals diversify within avian lineages, considering the constraints on color, location and processing of visual signals? The visual signals of some species are known to exhibit complementary colors, which have little spectral overlap, and tend to excite distinct sets of photoreceptors. **Complementary colors** represent a continuum lying on opposite sides of a line passing through the achromatic point (where all color receptors are equally stimulated) in color space (Endler 1992; Endler et al. 2005; Ham and Osorio 2007). The extremes of this continuum represent the most complementary colors, offering high contrast and discriminability when combined together, particularly over adjacent body regions (Endler 1992; Osorio et al. 1999a; Endler and Mielke 2005; Hill and McGraw 2006a). For example, in forest canopy birds, dwelling against a primarily green background, blue colors serve to increase the contrast of red colors against the background (Endler 1992). If each species within a lineage possesses a similar pair of complementary colors (as defined by the distribution in color space mentioned above), diverse patterns may evolve by redistributing these colors over the body. Thus, diverse yet high-contrast visual signals may evolve, supporting species discrimination. In addition to chromatic signals, luminance signals (achromatic or black- and-white variance) are also important to consider as they offer high contrast(Marchetti 1993; Mennill et al. 2003; Griggio et al. 2011), and may be prioritized for discrimination in certain circumstances(Schaefer et al. 2006). Thus, if we quantify color (and luminance) across all species within a lineage, we would predict 1) that plumage colors within a lineage would lie along a line in perceptual tetrachromatic color space (Goldsmith 1990; Endler and Mielke 2005), and 2) each species should possess colors that lie on opposite sides of the achromatic point within this continuum of complementary colors (Endler 1992) (Figure 1). As a corollary to this second, the presence of both complementary colors across species could alternatively indicate phylogenetic constraints due to shared ancestry. Therefore, if we perform phylogenetic comparative analyses of color and luminance scores (to account for phylogenetic non-independence), we also predict that we should uncover evidence of a departure from phylogenetic Brownian motion models of trait evolution over at least some regions of the body. This, together with the previous predictions, would be consistent with complementary colors being redistributed across body regions during diversification.

**Figure 1:**
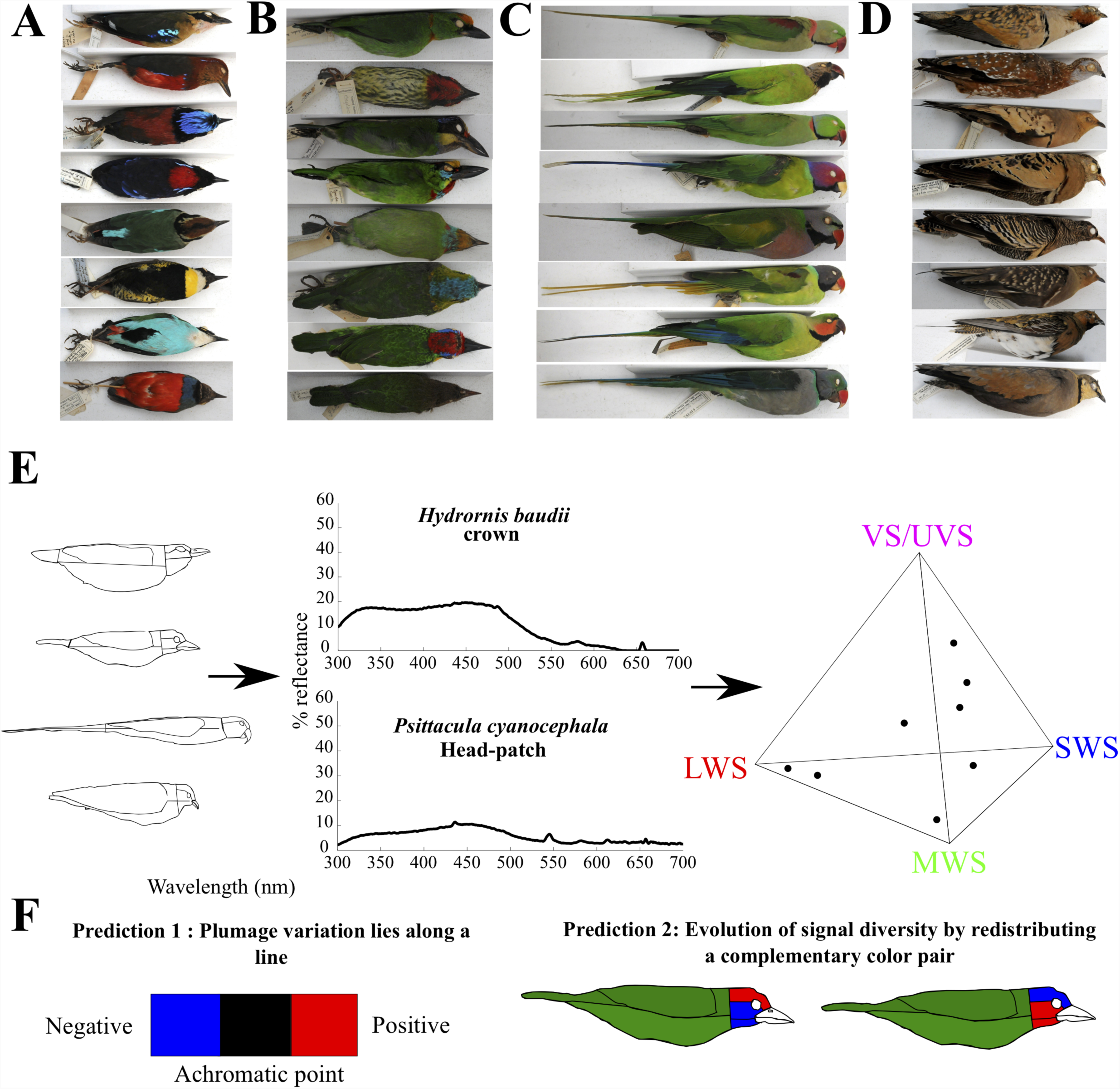
(A-D) Representative museum specimens of the four bird lineages examined in this study, the pittas (A), barbets (B), parakeets (C), and sandgrouse (D), from the collections of the Division of Birds, Smithsonian National Museum of Natural History, Washington D.C. (E) Workflow of analyses. Using museum specimens from all four lineages (left; the regions of the body are demarcated by lines), we measured reflectance spectra (examples in centre), and analyzed them using theoretical models of avian color vision including Goldsmith’s tetrahedron (right), where each vertex represents maximal relative excitation of one of the four cones (and therefore saturated colors). (F) Diagrammatic depiction of predictions. The color schemes here are illustrative, and are not meant to represent the real bird. We predict that plumage colors lie along a line representing putative complementary colors, which lie on opposite sides of the achromatic point on this line. We additionally predict that each species incorporates complementary colors for maximum contrast, and hypothesize that signal diversity evolves by redistributing a complementary color pair over the body.

Here, we describe the color space and interspecific color variation in four ecologically and phylogenetically diverse bird lineages, using ultraviolet-visible light reflectance spectrometry (Hill and McGraw 2006a). They are: 1) Pittas (Pittidae), understory invertebrate-eaters occurring from Africa to Australasia(Erritzoe and Erritzoe 1998), 2) Asian barbets (Megalaimidae), tropical forest-canopy frugivores(Short and Horne 2001), 3) Afro-Asiatic *Psittacula* parakeets (Psittacidae), fruit and seed-eaters inhabiting deciduous forests and woodland(Forshaw and Cooper 1989), and 4) Sandgrouse (Pteroclidae), arid-country ground-dwelling granivores (Maclean 1996) (Figure 1A-E, Supplementary Appendix). These families putatively represent both UVS- and VS-type avian visual systems (see Supplementary Figure 1 and Appendix for further discussion) (Stoddard and Prum 2011; Ödeen and Håstad 2013). We first test the prediction that plumage colors in each lineage largely lie along a single line in color space (Prediction 1). Secondly, we examine interspecific patterns in color or luminance scores across body regions and species, to address whether this linear distribution represents complementary colors that are redistributed over the body during diversification (Prediction 2) (Figure 1F). By identifying common interspecific patterns across these families with diverse life-histories and habitats, our study aims for insight into general processes underlying the evolution and perception of visual signals.

## Materials and Methods

### Museum specimens

We measured museum specimens of four avian lineages (Number of specimens, Number of species measured: Pittas: 80,28; Barbets: 81,30; Parakeets: 55,12; Sandgrouse: 57,16)(del Hoyo et al. 2014), held in the collections of the Division of Birds, Smithsonian National Museum of Natural History (USNM), Washington, D.C., USA (total 273 specimens, Supplementary Dataset). Where possible, we measured specimens collected relatively recently (Armenta et al. 2008), male and female specimens of sexually dichromatic species, and distinct subspecies (also see Supplementary Appendix) to obtain a comprehensive estimate of the color space occupied by each family.

### Reflectance spectrometry and photon catch of color cones

We measured plumage reflectance of museum skins, using an S2000 UV-visible fiber-optic reflectance spectrophotometer (Ocean Optics, Inc.) with a DT1000 deuterium-tungsten halogen light source. Measurements were referenced to a CIE D65 (white under average daylight illumination) white standard (Milton Roy Color Products), and dark referenced to a black surface. We first moved the probe over each region of the body, looking at the computer display to ensure that (to the best of our ability) we did not miss any patches that are not visible to the human eye (particularly cryptic UV sexual dimorphism). We then measured one reflectance spectrum for each color patch on each specimen using the Overture software. Although our dataset did not take into account within-patch variation as a result, we generally observed that intraspecific variation (and qualitatively, within-patch variation observed by moving around the probe) for the same patch was lower than interspecific variation, and is thus unlikely to alter the patterns we observe.

We used the MATLAB (MathWorks, Inc.) program TetraColorSpace(Stoddard and Prum 2008) and the R (R Core Team 2013) package PAVO(Maia et al. 2013) to analyze reflectance spectra. These algorithms incorporate cone sensitivities for averaged VS and UVS avian visual systems, to calculate theoretical photon catch for each cone (this representing the signaler phenotype, or visual signal under idealized light conditions (Stoddard and Prum 2008). Although the use of averaged visual systems does not directly model perception for each species, photon catch provides an objective way to quantify spectral signal in different portions of the avian-visible spectrum(Burkhardt 1989; Goldsmith 1990; Endler and Mielke 2005). We calculated photon catch of the four color cones using both programs (the values were concordant across both), performing the von Kries correction (Vorobyev and Osorio 1998) using a uniform white light (or idealized light) spectrum. Birds process luminance information separately from color information(Vorobyev and Osorio 1998; Endler and Mielke 2005), using the double cones(Goldsmith and Butler 2005). Thus, we also used PAVO to calculate the photon catch of the double cones as a measure of luminance, using known sensitivities for the double cone of the blue tit (*Cyanistes caeruleus*)(Hart et al. 2000). Again, although this does not directly represent luminance perception by each species, it provides an objective comparison of luminance differences in plumage. Using the relative photon catch values for each cone, we visualized plumage colors of each bird family in Goldsmith’s tetrahedral color space(Burkhardt 1989; Goldsmith 1990).

### Analyses

After obtaining raw photon catch values for each cone, we transformed these values into a three-dimensional XYZ color space representing the receptor-noise limited model of tetrachromatic color vision(Vorobyev and Osorio 1998; Vorobyev et al. 1998; Vorobyev 2003; Siddiqi et al. 2004). This was accomplished using the Weber fraction of each cone, which is calculated using the signal:noise ratio and the relative abundance of each cone in the retina. We incorporated published Weber fractions of the four cones for *Leiothrix lutea*(Vorobyev et al. 1998) as described in the literature(Cassey et al. 2008; Delhey et al. 2015), to transform photon catch values for each color patch into XYZ coordinates using a custom-written MATLAB code. The advantage of this color space is that distances between points are expressed in just noticeable differences (JND), an indication of the perceptual distance between them (Vorobyev and Osorio 1998; Siddiqi et al. 2004; Cassey et al. 2008; Pike 2012), thus providing a better approximation of how differences in color are perceived by the avian visual system. We also plotted color distributions for each family in this color space using the RGL package(Adler et al. 2003) in R sensu (Delhey et al. 2015).

To test Prediction 1, that plumage colors should largely distribute along a single line(Endler et al. 2005), we estimated the proportion of variation in coordinate space explained by the first major axis using principal components analysis (PCA) on the XYZ coordinates obtained above, following published studies(Cassey et al. 2008; Delhey et al. 2015). In order to test prediction 2, that this line represents a complementary color axis, and that the positions of these colors may be redistributed across the body during the diversification of these avian lineages, we required a metric that included not only the distance of each color from the origin (indicating “how complementary” a color is along the continuum), but which distinguished colors lying on opposite sides of the achromatic point (information which is lost in Euclidean distance measures). To achieve this, we transformed the XYZ coordinates into a spherical coordinate space in MATLAB, with the achromatic point at the origin. We used this coordinate (in radians) as a “color score” in subsequent analyses (using a species average, also see Results). By the definition of a complementary color pair as detailed above, they should therefore exhibit color scores with opposite signs. This is because they occur on opposite sides of the achromatic point(Endler 1992; Endler and Mielke 2005), and exhibit little spectral overlap (Ham and Osorio 2007). Additionally, this enabled us to transform complex measurements of color space into a ‘trait’ that could be compared using comparative phylogenetic analyses, while simultaneously testing hypotheses about the evolution of complementary colors. To examine whether each species possessed both colors in a complementary color pair, we constructed histograms of the maximum and minimum color score for each species within a family. Finally, we used phylogenetic comparative analyses to investigate whether color and luminance scores across each body region exhibited phylogenetic signal. We first sorted all the patches measured in each of the four avian lineages into crown, cheek, throat, back, wing, tail, and underpart patches (except the parakeets, where we measured crown, cheek, back, wings, underparts and both upper and undertail, see Supplementary Appendix). Next, we calculated the average color score and luminance index (double cone photon catch) for each region of the body for the male plumage of each species (to account for some species possessing more color patches than others, and thus enable direct comparisons). Using published phylogenetic information for each family (Groombridge et al. 2004; Irestedt et al. 2006; Jetz et al. 2012; Kundu et al. 2012; Den Tex and Leonard 2013) and the ape and phytools packages (Paradis et al. 2004; Revell 2012) in R, we calculated Pagel’s λ, a measure of phylogenetic signal, for color and luminance scores of each body region. This index measures whether trait evolution (in this case, color and luminance scores) follows a Brownian motion model of evolution, where phylogenetic effects drive trait evolution. In this scenario, the λ value is 1, whereas departures from Brownian motion result in a value lower than 1 (Pagel 1999; Münkemüller et al. 2012). To estimate the significance of the measured statistic, we compared this value to 1000 randomized values obtained using the inbuilt functions of the phytools package. To further verify these results, we additionally performed a second analysis. Using a phylogenetic distance matrix derived from the ape package, we calculated Mantel correlations between this matrix and an interspecific trait distance matrix derived for color and luminance for each body region (see Supplementary Data). This test provided additional quantification on the effects of phylogenetic relatedness on interspecific color variation.

## Results

### Prediction 1: Plumage colors lie along a line in tetrachromatic color space

Across the four avian lineages, we find that plumage colors distribute between two points in tetrahedral color space. The color signals of pittas lie between red (LWS) and violet (VS) color vertices (indicating highly-saturated colors) (Figure 2A). Barbets largely distribute between the green (MWS) and red (LWS) vertices, with a few blue-violet patches (Figure 2C). Plumage colors of *Psittacula* parakeets lie between the middle of red-green space and the middle of blue-uv space (Figure 2E), with a few patches near the LWS and MWS vertices. Finally, the plumage colors of sandgrouse are restricted to a region between the black achromatic point (the centroid)(Stoddard and Prum 2008) and the LWS (red) vertex (Figure 2G). The XYZ color space using the noise-limited model also recovers a linear axis of color variation, suggesting that this “axis” is genuine, and not an artifact of the tetrahedral color space. The results of PCA to quantify the proportion of variation explained by this line are summarized below for each avian lineage, and also in Supplementary Table 1 (see Supplementary Dataset):

**Table 1:** Pagel’s λ, a measure of phylogenetic signal (i.e. fit to a Brownian motion model of trait evolution) for different body regions for each of the four bird lineages. Values given represent λ for color scores (top) and luminance below. Values that are in bold are significant at a P-value of 0.05 (P-value obtained by comparison to 1000 randomized phylogenetic trees for each region). For each family, below the λ value are two rows indicating the means and coefficient of variation (as a percentage) for each body region for both color and luminance scores.

| Color | Crown | Cheek | Throat | Back | Wing | Tail<br>(Upper<br>tail) | Undersides | Under<br>tail |
| --- | --- | --- | --- | --- | --- | --- | --- | --- |
| **Pittas ($\lambda$)** | 0.00007 | **0.999** | 0.28 | 0.14 | **0.84** | **0.999** | 0.00006 |  |
| **Mean** | 0.3 | 0.26 | 0.51 | 0.14 | -0.06 | -0.21 | 0.49 |  |
| **CV(%)** | 156.22 | 114.20 | 58.40 | 445.51 | 1286.41 | 347.03 | 75.51 |  |
| **Asian**<br>**Barbets ($\lambda$)** | **0.999** | **0.61** | **0.57** | **0.999** | **0.72** | **0.82** | **0.91** |  |
| **Mean** | 0.52 | 0.13 | 0.26 | 0.53 | 0.23 | 0.06 | 0.44 |  |
| **CV(%)** | 90.41 | 415.05 | 206.99 | 41.74 | 139.16 | 523.32 | 28.06 |  |
| **Afro-**<br>**Asiatic**<br>**Parakeets**<br>**($\lambda$)** | 0.00006 | 0.00006 |  | 0.39 | 0.26 | 0.00006 | 0.00006 | 0.17 |
| **Mean** | 0.36 | 0.38 |  | 0.54 | 0.03 | -0.45 | 0.54 | 0.79 |
| **CV(%)** | 74.61 | 65.74 |  | 35.24 | 2263.16 | 105.04 | 13.22 | 5.10 |
| **Sandgrouse**<br>**($\lambda$)** | 0.04 | 0.00007 | 0.00007 | 0.00007 | 0.00007 | 0.00007 | 0.00007 |  |
| **Mean** | 0.67 | 0.60 | 0.50 | 0.83 | 0.75 | 0.66 | 0.69 |  |
| **CV(%)** | 28.35 | 20.41 | 42.82 | 12.01 | 17.39 | 31.70 | 25.78 |  |
| **Luminance** |  |  |  |  |  |  |  |  |
| **Pittas ($\lambda$)** | **0.88** | 0.00006 | 0.00006 | 0.25 | **0.71** | 0.38 | 0.00006 |  |
| **Mean** | 0.03 | 0.02 | 0.10 | 0.06 | 0.09 | 0.03 | 0.07 |  |
| **CV(%)** | 113.93 | 156.62 | 79.46 | 101.64 | 84.59 | 117.61 | 61.69 |  |
| **Asian**<br>**Barbets ($\lambda$)** | **0.88** | **0.38** | **0.68** | **0.92** | 0.00006 | 0.00006 | **0.92** |  |
| **Mean** | 0.03 | 0.05 | 0.06 | 0.01 | 0.01 | 0.01 | 0.08 |  |
| **CV(%)** | 76.28 | 63.50 | 54.78 | 60.56 | 69.97 | 63.53 | 62.25 |  |
| **Afro-**<br>**Asiatic**<br>**Parakeets**<br>**($\lambda$)** | 0.00006 | 0.13 |  | 0.00006 | 0.087 | 0.00006 | 0.00006 | 0.00006 |
| **Mean** | 0.11 | 0.08 |  | 0.09 | 0.04 | 0.08 | 0.11 | 0.13 |
| **CV(%)** | 38.89 | 60.08 |  | 65.06 | 43.77 | 48.45 | 40.64 | 36.59 |
| **Sandgrouse**<br>**($\lambda$)** | 0.21 | 0.00007 | 0.00005 | 0.39 | 0.82 | 0.00007 | 0.12 |  |
| **Mean** | 0.07 | 0.13 | 0.12 | 0.03 | 0.06 | 0.06 | 0.06 |  |
| **CV(%)** | 69.08 | 56.07 | 65.18 | 123.17 | 83.93 | 115.84 | 93.06 |  |

**Figure 2:**
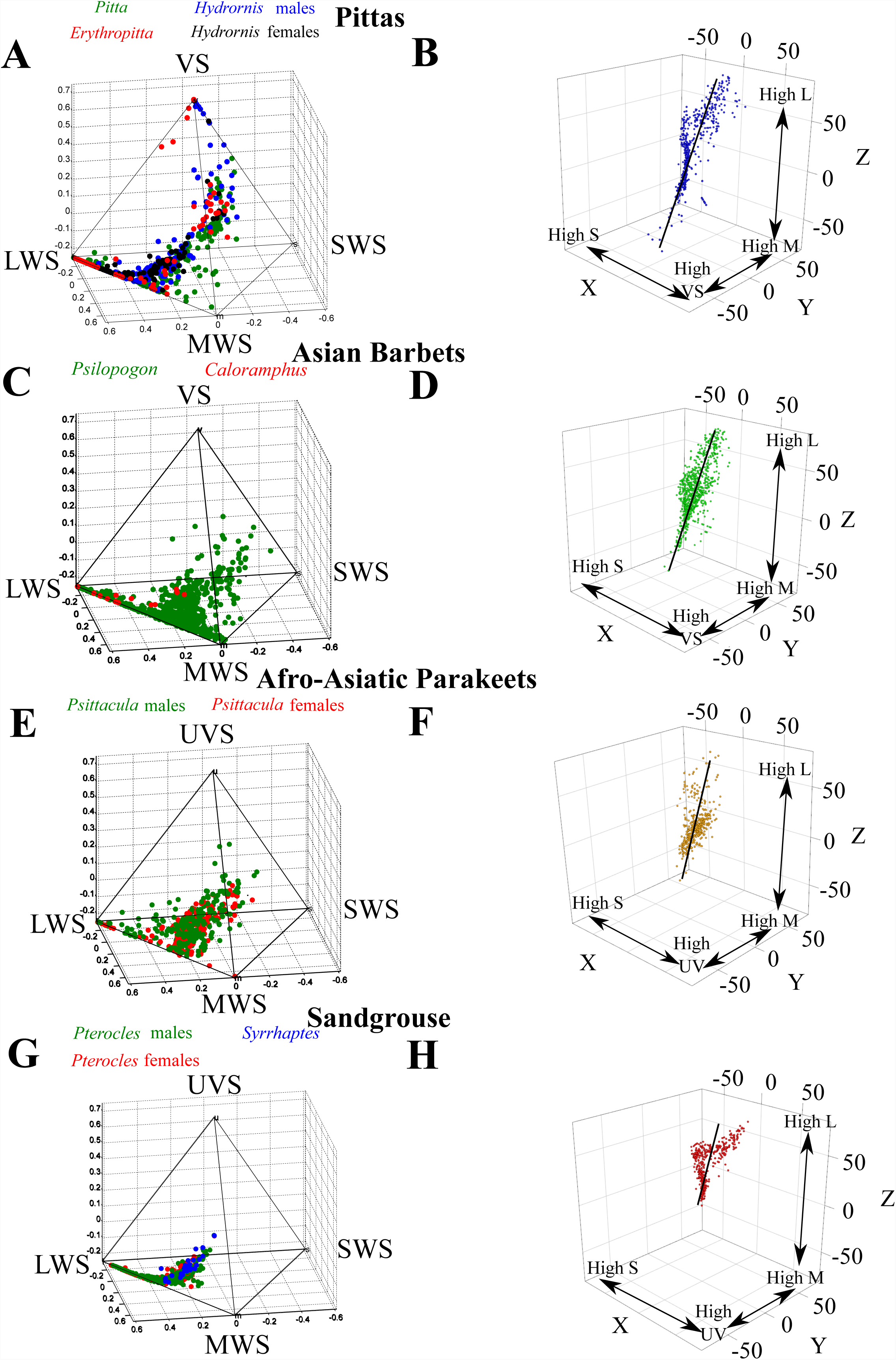
Color space occupancy and analyses of signal variation in pittas (A,B), barbets (C,D), parakeets (E,F) and sandgrouse (G,H). Left-hand side figures represent color space occupied by each family (1 point/color patch measured), as visualized using Goldsmith’s tetrahedron. Each vertex represents relative photon catch of a particular cone (see Figure 1). Right hand side plots represent the same data points transformed into a three-dimensional XYZ color space using a noise-limited model of avian tetrachromatic vision. The black lines through the points represent the first major axis (PC1) of chromatic variation.

#### Pittas

PC1 (the major axis of variation) of the XYZ coordinates in color space explains 85% of chromatic variation (Figure 2B). PC1 loads weakly negatively on X (-0.15), and exhibits strong positive loadings (0.6 and 0.78) on Y and Z, respectively.

#### Barbets

PC1 explains almost 74% of chromatic variation (Figure 2D), loading weakly negatively on X (-0.03), moderately positively on Y (0.465), and strongly positively on Z (0.884).

#### Parakeets

PC1 explains 75% of variation in color (Figure 2F), loading weakly negatively on X (-0.2), moderately positively on Y (0.57) and strongly positively on Z (0.8).

#### Sandgrouse

PC1 explains about 70% of color variation, loading weakly negatively on X (-0.32) and strongly positively on Y and Z (0.64 and 0.7).

Across all four lineages, the Z coordinate loads most strongly on PC1, thus suggesting that most variation in perceptual coordinate space occurs along the elevational rather than azimuthal direction along the PC1 line. Therefore, in subsequent analyses, we used the elevational coordinate Φ and the sign of this coordinate as an indicator of where different colors lie along this line. Although this does not take variation in the azimuthal plane into account, the results of our analysis suggest that this variation is negligible compared to variation along the elevational axis in all four families. Thus, colors with opposite signs of Φ in this dataset lie on opposite sides of the achromatic point (as is evident from the spread of the data in Figure 2). We used PCA only to estimate the proportion of variance along this line, and not in any subsequent analysis.

### Prediction 2: Color space axes represent complementary colors, lying on opposite sides of the achromatic point

We next test if color pattern diversity is achieved by redistributing complementary colors across certain body regions using species averages for color and luminance scores across body regions. We predict a) that species within families exhibit colors lying on both sides of the achromatic point, and b) that measures of phylogenetic signal indicate departure from a Brownian motion model of evolution, consistent with changing positions of a complementary color pair during diversification. We summarize the results of these analyses below:

#### Pittas

After transforming into a spherical coordinate space, elevation coordinates span between -1.54 and +1.57 across the family, i.e. on opposite sides of the achromatic point and at roughly equal distances from it along the elevation axis, consistent with the interpretation of a complementary color axis. For example, the deep-blue (to human eyes) crown of the male *Hydrornis baudii* has, on average, a color score of -1.15, and the deep-red crown of the sympatric(Erritzoe and Erritzoe 1998) *Erythropitta granatina* scores +1.12. This is also consistent with a hypothesis of complementary colors, in that these colors also represent opposite ends of the avian-visible light spectrum. Histograms of maximum and minimum color scores of each species within the family show that most of these species possess both complementary colors, the peaks of these distributions lying on opposite sides of the achromatic point (Figure 3A). Phylogenetic comparative analyses (Table 1) reveal that patterns of plumage evolution are heterogeneous across the body regions of pittas. Color scores are consistent with a Brownian motion model of evolution on the cheek, wing and tail, and exhibit weak and non-significant phylogenetic signal across other body regions. Luminance scores exhibit significant phylogenetic signal only on the crown and wing. Mantel tests for correlation between phylogenetic and trait distance broadly corroborate these results: luminance distance correlates significantly with phylogenetic distance only on the wing, whereas color correlates on the cheek, wing and tail (Supplementary Data). In addition, the regions with non-significant phylogenetic signal all possess relatively high coefficients of variation in color scores (Table 1). Thus, plumage patterns in the pittas are consistent with species possessing colors on opposite sides of a continuum of complementary colors, and with the body distributions of these colors being swapped across certain body regions during diversification.

**Figure 3:**
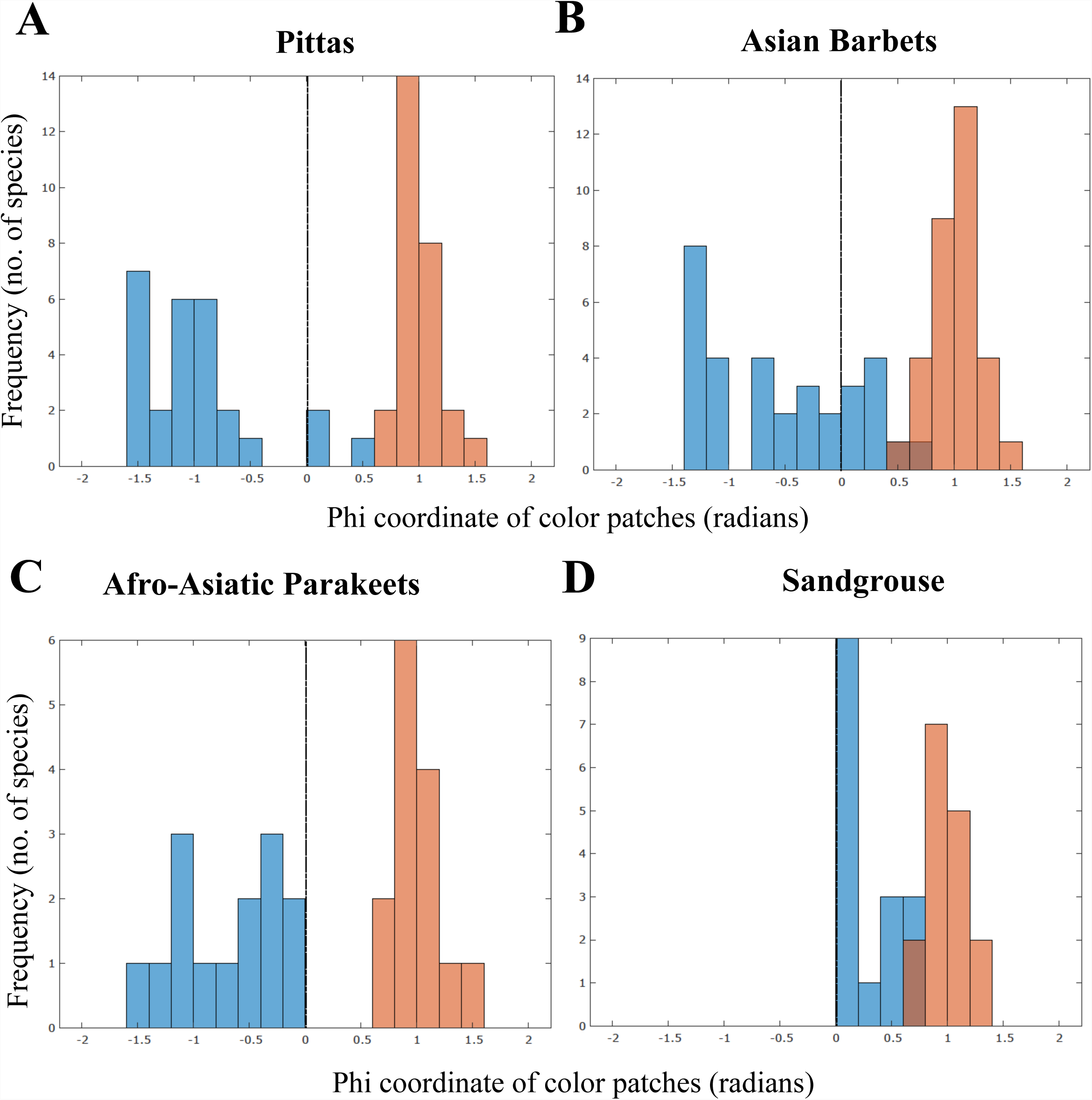
Linear axes of plumage variation represent complementary colors. Shown here are histogram distributions of maximum (red) and minimum (blue) color scores (the maximum and minimum phi-coordinate in radians) of each species within four bird lineages, the pittas (A), Asian barbets (B), Afro-Asiatic parakeets (C), and sandgrouse (D). In the first three families, most species possess colors that are on opposite extremes of the continuum of complementary colors, with maximum and minimum scores on opposite side of the achromatic point (zero). In the fourth, the sandgrouse, colors are clustered on one side of the achromatic point, and thus do not exhibit chromatic complementarity.

#### Barbets

Color scores span between -1.4 and +1.57, also consistent with a complementary color axis. For example, the red throat of *Psilopogon mystacophanos*, exhibits, on average, a color score of +1.1, and the turquoise throat of the sympatric(Short and Horne 2001) *P. rafflesii* a score of -1.21, which, like pitta colors, lie on opposite sides of the achromatic point and on a line through it. Again, histograms of color distribution (Figure 3B) demonstrate that most barbet species exhibit colors lying on opposite sides of the achromatic point (i.e. complementary colors). Color and luminance scores (Table 1) exhibit significant λ values across all regions (except luminance scores on the wing and tail), but values for head patches (particularly the cheek and the throat) are much lower than 1 (0.61 and 0.57), indicating a departure from a Brownian motion model of trait evolution. Phylogenetic and color distance are correlated on all body regions, but not on the head regions (Supplementary Data), corroborating the results from phylogenetic signal. In addition, all head regions possess relatively high CVs for color scores, but not body regions (except the wing, which does, however, exhibit phylogenetic signal suggesting that this variation has a phylogenetic component). Taken together, these results are also consistent with body colors being a constrained feature within this lineage, but colors being swapped around on the head.

#### Parakeets

Color scores span between -1.47 and +1.57, again consistent with a complementary color axis. For example, the wing of the male *Psittacula longicauda nicobarica* (-1.38) exhibits the opposite sign to the red shoulder patch of male *P. cyanocephala* (+1.32). Color histograms again indicate the presence of both complementary colors on all species within the family, with maximum and minimum scores for each species lying on opposite sides of the achromatic point (Figure 3C). Neither color nor luminance scores exhibit significant phylogenetic signal across any body regions (Table 1) when compared to a Brownian motion model of trait evolution, and additionally do not exhibit significant correlations with phylogenetic distance (Supplementary Data). Color scores exhibit higher CVs than luminance scores (Table 1), which is also consistent with signal diversification along a chromatic complementary axis.

#### Sandgrouse

Color scores span between 0 (the achromatic point) and +1.33. This suggests that sandgrouse are clustered in chromatic space to one side of the achromatic point, further supported by color histograms (Figure 3D). However, aside from luminance scores on the wing (Table 1), neither color nor luminance scores exhibit significant phylogenetic signal on any body region. In addition, other than color scores on the crown (Supplementary Data), interspecific color and luminance distances are not significantly correlated with phylogenetic distance. Coefficients of variation of color scores (Table 1) are generally lower than those for luminance across body regions, unlike the other three lineages. Taken together with the apparent lack of chromatic complementarity in sandgrouse, interspecific luminance variation may putatively play a greater role in signal diversification. The sympatric *Pterocles alchata* and *P.orientalis*(Benítez-López et al. 2014) represent a noteworthy example of such divergence. The black belly of the male *P.orientalis* exhibits an average luminance score <0.0001, whereas the white belly of the male *P.alchata* exhibits an average luminance score of 0.34.

## Discussion

Across families, we thus uncover consistent evidence that complementary colors (or putatively patches of different luminance in sandgrouse, representing black-and-white variation, which are also inverses of each other) underlie the evolution of contrasting visual signals. Most species within each lineage possess both colors of a complementary pair, and phylogenetic comparative analyses indicate generally low or insignificant phylogenetic signal (except the barbets, where, however, the head regions diverge from a Brownian motion model of trait evolution). Taken together with the constrained distribution of color scores (largely along a single line in color space), we hypothesize that evolutionary diversification of color patterns occurs by redistributing a complementary color pair across body regions. We discuss this further below.

### Visual signals and complementary colors

To summarize, we find that plumage colors in each of the four bird lineages distribute along an axis between two colors (or regions of the avian-visible spectrum) which are complementary (spanning either side of the achromatic point), except the arid-country sandgrouse whose colors are found to only one side of the achromatic point. Additionally, based on phylogenetic comparative analyses, we hypothesize that signals diversify by redistribution of complementary colors. In the case of sandgrouse, the lack of phylogenetic signal and chromatic complementarity, together with the larger variance in luminance scores across body regions compared to color scores, leads us to tentatively hypothesize that signal evolution in this family has occurred along an achromatic (black-white) complementary axis rather than a chromatic one. Although not directly confirmed in our study systems, it is important to note here that tetrachromatic visual systems (avian and reptilian) possess a number of opponent color processes to compare cone outputs(Yazulla and Granda 1973; Osorio et al. 1999b; Ventura et al. 2001; Smith et al. 2002; Goldsmith and Butler 2005; Rocha et al. 2008). In human trichromatic visual systems, red-green, yellow-blue and luminance (black-white) opponent comparisons result in all perceived hues occupying a continuum between these perceptually distinct opponent colors(Hurvich and Jameson 1957). Different opponent mechanisms (or color axes) dominate at various wavelengths and intensities of ambient light, accordingly shifting the perceived color space (the Bezold–Brücke phenomenon)(Boynton and Gordon 1965). This may represent a putative mechanism enabling discrimination of complementary colors across light environments, although we note that we do not possess the evidence at present (i.e. physiological data) to explicitly test this.

We uncover evidence both that most species within a lineage possess complementary colors in their plumage (Figure 3), and also that phylogenetic patterns of trait evolution depart from Brownian motion over at least some body regions in all families (Table 1). Taken together, this is consistent with complementary colors in their plumage, putatively for high chromatic contrast (Endler 1992) under constraints, and with signal diversification by redistributing these complementary colors over certain regions of the body. In the case of pittas, these regions appear to be the crown, throat, back and underparts. For example, *Hydrornis baudii* possesses a blue crown and underparts, and a reddish-brown back, whereas the sympatric *Erythropitta granatina* possesses a deep blue-violet back and a bright red crown patch and belly. For Asian barbets, this redistribution of colors appears to occur primarily on the cheek and throat, and an examination of their color patterns supports this. Most members of the family possess largely green bodies, and bright colors are confined to the head regions. For parakeets, a lack of phylogenetic signal across the body suggests that diversification may occur by redistribution of colors across all body regions. This is consistent with the fact that different species possess both short-and long-wavelength colors on the head, wing, underparts and tail. Within all four lineages, multiple species occur in sympatric assemblages(Forshaw and Cooper 1989; Maclean 1996; Erritzoe and Erritzoe 1998; Short and Horne 2001; Groombridge et al. 2004; del Hoyo et al. 2014; Krishnan and Tamma 2016). Complementary colors may thus support pattern discrimination between species (or sexes of dichromatic species)(Hill and McGraw 2006b; Osorio and Vorobyev 2008). Future studies using image analysis techniques (such as those used to study egg patterns)(Stoddard et al. 2014), will aim to quantify and obtain further insight into how patterns of complementary colors have evolved.

### Ecological processes may constrain plumage diversity to complementary color axes

How do ecological constraints influence plumage colors, and how might these explain patterns observed in our study? Lineages descended from a common ancestor may be predicted to resemble each other; alternatively, sexual selection and species recognition may accelerate signal diversification (Seddon et al. 2013; Mason et al. 2014). Signals may also be constrained by ecological factors to a specific complementary color axis. For example, the red colors of pittas and barbets are due to carotenoids(Thomas et al. 2014), putatively derived from dietary sources(Hill et al. 2002), in contrast to structural short-wavelength colors (Saranathan et al. 2012). Parakeet pigment colors are due to psittacofulvins (McGraw and Nogare 2004). Finally, sandgrouse do not possess plumage carotenoids(Thomas et al. 2014), and pigmentation is thus likely to be primarily melanin-based (brown-black). This may constrain plumage diversification to an achromatic axis (or to changes in barring and speckling, which our study did not investigate), albeit with the caveat that luminance variation is difficult to compare using museum specimens. However, a comparison of plumage patterns in sandgrouse (Figure 1) reveals that many species possess conspicuous black and white patches, whose distributions differ between species. Some possess these patches on the face, others on the wings and belly. Similar patterns of evolution along an achromatic axis may have also putatively occurred in other melanin-pigmented bird groups, such as larks, bustards, and coursers, as well as many raptors(del Hoyo et al. 2014), and merit further investigation.

Ecological pressures of sensory drive (for example, crypsis from predators and conspicuousness to intended receivers) may additionally constrain plumage diversity in bird lineages where all members exhibit relatively similar ecological preferences. All four families studied here experience predation, and possess both cryptic colors, and colors that offer maximal contrast in their preferred habitats. For example, blue-violet and saturated reds are very conspicuous against a forest understory background(Siddiqi et al. 2004), and reds also against the green forest canopy, where blue serves to increase within-pattern contrast(Endler 1992; Gomez and Théry 2007); these are the colors exhibited by pittas and barbets, which typically occupy these habitats(Erritzoe and Erritzoe 1998; Short and Horne 2001) (Figure 2). Cryptic colors, defined as matching the background in a habitat(Endler 1992; Gomez and Théry 2007) (green in tree-dwelling barbets and parakeets, reddish-brown in ground-dwelling pittas and sandgrouse), also occur across all four families, which are additionally noted in the literature as being unobtrusive, camouflaged or difficult to locate within their habitats (see discussion in Supplementary Online Appendix). It is possible that the composition of ambient light exerts constraints on plumage colors by shifting the perceived color space (Boynton and Gordon 1965; Wright 1975), although birds are known to exhibit color constancy, and the effects of this shift may be minor (Olsson et al. 2016). Alternatively, microhabitat variation in light composition may influence which colors are the most conspicuous (Endler and Thery 1996; Uy and Endler 2004), as well as whether birds use chromatic or achromatic contrasts in pattern discrimination (Endler and Thery 1996; Schaefer et al. 2006). However, we have not directly measured the light environments inhabited by these species. Additionally, our use of theoretical models makes the assumption that all species within a lineage perceive color the same way, whereas some differences between species are likely to exist. The effects of sensory drive must thus be treated as a tentative hypothesis, and field data are needed to further understand both the predation these birds experience, and the light microhabitats they use.

Color vision is challenging to study comparatively in speciose bird lineages containing rare or range-restricted species, and many of the species we examine are poorly known. Thus, although our study design does not permit us to conclusively identify the ecological driver of these patterns, we do find consistent evidence of overall patterns of evolution along complementary color (or achromatic) axes in a manner that does not depend on their phylogenetic relatedness. Based on this, we hypothesize that color patterns may have diversified by redistribution or replacement of these complementary colors between species. We speculate that this mechanism supports the evolution of highly contrasting, yet diverse recognition signals under constraints (dietary and sensory, among others), and further research will focus on the ecological processes underlying these constraints. Bird lineages that have undergone multiple niche shifts would provide a suitable crux to address ecology and plumage diversification.

## Acknowledgements

We are indebted to Helen James for providing access to the reflectance spectrometer and collections at the Smithsonian museum, and for helpful discussions and feedback on the data and manuscript. Preliminary work on this project was carried out when AK was a postdoctoral fellow at Johns Hopkins University. We thank Brian Schmidt, Chris Milensky, Christina Gebhard and Gary Graves for help with the bird collections and access to instruments, Chad Eliason, Sushma Reddy, Sutirth Dey, Ramana Athreya, Teresa Feo, Terry Chesser, Uma Ramakrishnan, Wu-Jung Lee and Sanjay Sane for feedback and discussions.

## Authors’ Contributions

Conceived and designed the study: AK, Collected data: AK AS, Analyzed and interpreted the data: AK KT, Wrote the first draft of the manuscript: AK, Contributed to editing and revision of the manuscript: AK AS KT. All authors approved submission of the manuscript.

