## Supplementary Appendix for "Visual signal evolution along complementary color axes in four bird lineages"

**Supplementary Appendix: Visual signal evolution along complementary color axes in four bird radiations**

*Bird families: Taxonomy and visual systems*

The four bird radiations in this study are ecologically diverse, occurring in multispecies assemblages throughout their large ranges(del Hoyo et al. 2014). Below, we provide a brief description of each, together with a note on what is known about their visual systems(Stoddard and Prum 2011; Ödeen and Håstad 2013). The taxonomy we note here is based on BirdLife International (2016), although many of the species-level splits (the 12-way split of the Red-bellied Pitta raised the number of species from 34 to 46) had not been adopted when the study was carried out, and we have therefore also included the proportion of previously recognized species (BirdLife International 2014) total to illustrate our species coverage at the time of the study. Changes for non-passerines had been adopted earlier, and these changes therefore mainly affect the nomenclature of the Pittidae in our dataset; for other families, we include only one species total.

**Pittas**

Pittas (Pittidae; our dataset sampled 28/46 currently recognized species, or 27/34 previously recognized species) are suboscine terrestrial passerines that occupy forest understory habitats, and feed on insects, worms and other invertebrates(Erritzoe and Erritzoe 1998). Although all members of the family were previously treated as congeneric in the genus *Pitta*, recent treatments have recognized three distinct groups that are now separated as genera(del Hoyo et al. 2014). The genus *Hydrornis* are sexually dichromatic, and occur from the Eastern Himalayas through Southeast Asia to Java and Bali. The genus *Erythropitta* are found on the Malay Peninsula, Borneo, Sumatra, the Philippines, and through Australasia to Northern Australia. The genus *Pitta* as currently defined is the most geographically widespread, with several partially migratory species. Two species occur in tropical Africa, and others from the Indian Subcontinent to Northern Australia (two species), with endemic island forms in the Philippines, the Lesser Sundas, the Moluccas, and the Admiralty and Solomon Islands(Erritzoe and Erritzoe 1998). Pittas are closely related to broadbills, which are thought to possess a VS-type visual system, and we have therefore treated them as such as well(Stoddard and Prum 2011). We note, however, that modeling pitta plumage colors using a UVS system does not drastically alter their relative position in tetrahedral color space (slightly shifting them towards the blue axis) (Supplementary Figure 1), and is therefore unlikely to significantly alter the patterns we observe. Pittas exhibit unobtrusive behavior in the forest understory, and are very difficult to locate in spite of their striking plumage; predators include ground-dwelling mammals and birds of prey, although data are scarce(Erritzoe and Erritzoe 1998).

**Asian barbets**

35 species of Asian Barbets (Megalaimidae; our dataset sampled 30 of these) occur through the Asian tropics from the Indian Subcontinent to Java and Bali. Of these, 2 species are in the genus *Caloramphus*, which are social fruit-insect eaters of the mid-storey of Sundaic forests. The other 33 are now in the genus *Psilopogon* (32 of these were formerly placed in *Megalaima*), territorial canopy-level fruit eaters which nest in tree cavities and occur throughout their ranges in multispecies assemblages(Short and Horne 2001). Nearly all species occur in forest canopy habitats, although some also occur in deciduous habitats. Sexes are alike in nearly all species; only *P.mystacophanos* exhibits sexual dichromatism. Centres of endemism of *Psilopogon* barbets include Sri Lanka and Peninsular India (5 species total), the greater Sundas including Borneo, Sumatra, Java and Bali (14 species in total), and Vietnam, Laos, Thailand and Cambodia (6 species more or less restricted to this region). Analysis of opsin sequences suggest that barbets of this genus possess a VS-type visual system, and we have modeled them accordingly (see above)(Ödeen and Håstad 2013). Barbets are known to experience predation from birds-of-prey, and exhibit behaviors which render them difficult to locate in dense foliage(Short and Horne 2001).

**Afro-Asiatic Parakeets**

The third bird radiation we investigated is the long-tailed parakeets of the genus *Psittacula* (Psittacidae)*,* of which 12 species occur across South and Southeast Asia, with one of these (*P.krameri*) occurring in Africa as well. Restricted-range endemic species occur in Sri Lanka, the Western Ghats of India and the Southern Nicobar Islands. In addition to these, endemic species once occurred in the Mascarenes and the Seychelles, of which only one species (the endangered *P.eques*) survives today(Forshaw and Cooper 1989). We sampled all 12 mainland Asian species. Additionally, the endemic subspecies of *P.longicauda* from the Andaman (*P.l.tytleri*) and Nicobar (*P.l.nicobarica*) Islands are highly distinctive; our study included both of these in addition to the nominate *longicauda* in order to encompass as much phenotypic diversity within the genus as possible. All species in the genus exhibit a degree of sexual dichromatism. These parakeets are largely birds of woodland and deciduous forest (with a few species also occurring in evergreen habitats), where they feed on fruits and seeds. Multiple species may occur together, forming mixed flocks. We have modeled their plumage colors using a UVS-type visual system(Stoddard and Prum 2011). Predation pressure from falcons, owls and other raptors is known for several members of the parrot family, including these. Some members of the genus possess screeching calls that are issued at the approach of predators, and roost communally; however, they are said to be relatively quiet and inconspicuous when feeding in trees(Forshaw and Cooper 1989).

**Sandgrouse**

Sandgrouse (Pteroclidae), are sexually dichromatic, flocking granivores of arid, open habitats in Africa and Asia(Maclean 1996). A total of 16 species make up this family (our dataset sampled all 16), of which two (the genus *Syrrhaptes*) are birds of cold desert, steppe and alpine plateaux in central Asia(del Hoyo et al. 2014). Their ranges largely do not overlap with each other, and our analysis of interspecific variation and disparity focused on the other 14 species, the genus *Pterocles*. The habitats these 14 species occupy range from semi-arid grasslands and open country to stony and sandy deserts, and they fly long distances daily in search of water(Maclean 1996). Six species are found only in Sub-Saharan Africa, a seventh in Madagascar, another five from the Western Indian Subcontinent to Northern Africa through the Middle East (two of these are also found on the Iberian peninsula), and one species is endemic to the Indian Subcontinent. The last (*P.exustus*) is widely distributed from India and Pakistan through the Persian Gulf to the Sahel and East Africa(del Hoyo et al. 2014). Opsin sequences indicate the presence of a UVS system in sandgrouse(Ödeen and Håstad 2013), which is what we have used to model their plumage patterns. Sandgrouse are known to experience significant predation from both mammalian and avian predators, the latter including falcons, harriers and other raptors; an estimate suggests that a single raptor the size of a lanner falcon may consume about 300 sandgrouse a year(Maclean 1996).

*Specimen selection*

All specimens measured in this study are preserved in the collections of the Division of Birds, Smithsonian National Museum of Natural History, Washington D.C. We measured adult males and females of all species, wherever present in the collections. A particular concern with such studies is fading of specimens, which may alter plumage spectra. To offset this, we followed other studies in a) selecting specimens with no qualitative evidence of fading and b) where possible, measuring specimens of diverse collection ages(Armenta et al. 2008; Stoddard and Prum 2008; Doutrelant et al. 2016), including relatively recent specimens (recent implying the 1960s onwards until 2014). The Goldsmith model does not perform well on black patches (and may artificially inflate the relative photon catch of color channels, depending on specimen condition), and the TetraColorSpace program therefore treats these patches as possessing zero reflectance.

*Definitions of body regions for variance analyses*

Using photographs of each specimen (taken using a Nikon D300 DSLR camera in lateral, dorsal and ventral orientations), we divided the body of the bird into seven regions commonly used by ornithologists to describe plumage patterns: the crown (which we defined as all patches occupying the upper surface of the head to above the eye, extending backward to the nape), the cheek (defined as all patches occupying the region from the eye downwards to the base of the mandible, extending backward to the base of the neck), the throat (all patches below the base of the mandible), the upperparts (the back and rump), wing, tail (typically the upper surface, but see below), and underparts (breast, belly and vent)(Dale et al. 2015). For *Psittacula* parakeets, we did not treat the throat as a separate region from the cheek, owing to the structure of plumage patterns across their heads (briefly, the throat is a restricted region of overall head area in this genus, continuous with the black cheek stripes; these were included instead in our analyses of cheek patterns as they extend up the sides of the head) (refer to species plumages in (del Hoyo et al. 2014)). Secondly, owing to the long tails of these parakeets, we could obtain reliable measurements of undertail colors in all species, and these are therefore included in our analyses. Detailed plumage patterns, including speckling and mottling across families, will be analyzed in future studies.

*Further notes on plumage patterns*

A few plumage regions were not measured in our study. For reasons of accessibility due to the methods of skin preparation, we did not measure the underwing and undertail colors of any species (except the undertail of parakeets; see above), and we could not reliably access the white wing patches of certain Pittidae, which have also therefore been excluded from our analyses (these, however, are qualitatively similar across species, and likely serve a conspicuousness function during displays, but are not often visible in the perched bird)(Erritzoe and Erritzoe 1998). In addition, we could not access the tails of some species of pitta with the spectrometer (which were obscured by the wings). Finally, our analyses considered only plumage colors and not the colors of bare parts such as legs, bills or facial skin, owing to concerns about fading, specimen preservation and therefore the measurement of luminance. We do, however, include the bills of the *Psittacula* parakeets (largely bright red, with the exception of two species with orange bills, and black in the females of some species), in our tetrahedral plots of color space, but not in subsequent analyses (this is because specimen preparation may alter the brightness of color in some specimens). These bills are also likely contributors to the overall chromatic signal, although perhaps not to diversification between species.

Finally, although our dataset sampled almost all the extant diversity within these four families, a discussion of missing species in the dataset (that were not represented in the Smithsonian collections) is important to make here. We sampled all distinctive species of Asian barbets (most missing species are recently split subspecies, which exhibit relatively minor plumage differences that are unlikely to alter our findings) barring the Bornean *P.eximius*, which differs from its closest relatives both in its black throat and in possessing red patches on the crown and throat(Short, L.L and Horne 2001). Our dataset also did not include six species of pitta in the previously recognized taxonomic arrangement (and we examined two representative members of the recently split *E.erythrogaster* species complex, which, however, broadly resemble each other in global plumage patterns), which are also close relatives of species that were included, and possess qualitatively similar plumage patterns: *P.reichenowi* is similar to *P.angolensis* except for its dark green breast, *P.elegans* and *P.anerythra* broadly resemble *P.versicolor*, and *P.superba* resembles *P.maxima* except for its black and red (as opposed to white and red) underparts. Meanwhile, *E.venusta* and *E.kochi* closely resemble *E.ussheri/granatina* and *E.erythrogaster* respectively in global color patterns (although they are morphologically divergent). The addition of these species is thus unlikely to significantly alter overall patterns in the family. In the parakeets and sandgrouse, our coverage was nearly complete: we measured all extant parakeets except for the island *P.eques*, and all 16 sandgrouse species (del Hoyo et al. 2014).
