## Supplementary Figure 1 for "Visual signal evolution along complementary color axes in four bird lineages"

### Color Space Occupancy

#### *Erythropitta* and *Hydrornis* males combined

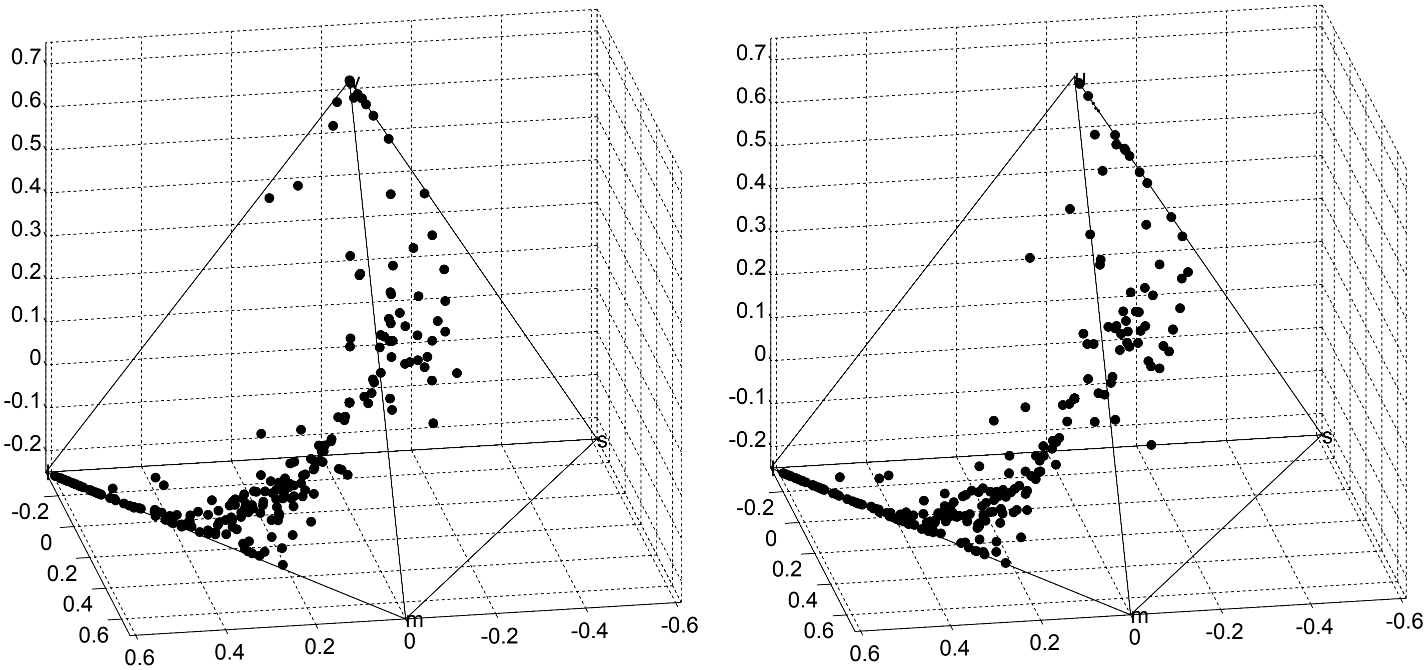


VS UVS

Figure S1: Comparison of color space occupancy in pittas modeled using VS and UVS visual systems
